# Naked mole-rat acid-sensing ion channel 3 forms nonfunctional homomers, but functional heteromers

**DOI:** 10.1101/166272

**Authors:** Laura-Nadine Schuhmacher, Gerard Callejo, Shyam Srivats, Ewan St. John Smith

## Abstract

Acid-sensing ion channels (ASICs) form both homotrimeric and heterotrimeric ion channels that are activated by extracellular protons and are involved in a wide range of physiological and pathophysiological processes, including pain and anxiety. ASIC proteins can form both homotrimeric and heterotrimeric ion channels. The ASIC3 subunit has been shown to be of particular importance in the peripheral nervous system with pharmacological and genetic manipulations demonstrating a role in pain. Naked mole-rats, despite having functional ASICs, are insensitive to acid as a noxious stimulus and show diminished avoidance of acidic fumes, ammonia and carbon dioxide. Here we cloned naked mole-rat ASIC3 (nmrASIC3) and used a cell surface biotinylation assay to demonstrate that it traffics to the plasma membrane, but using whole-cell patch-clamp electrophysiology we observed that nmrASIC3 is insensitive to both protons and the non-proton ASIC3 agonist 2-Guanidine-4-methylquinazoline (GMQ). However, in line with previous reports of ASIC3 mRNA expression in dorsal root ganglia (DRG) neurons, we found that the ASIC3 antagonist APETx2 reversibly inhibits ASIC-like currents in naked mole-rat DRG neurons. We further show that like the proton-insensitive ASIC2b and ASIC4, nmrASIC3 forms functional, proton sensitive heteromers with other ASIC subunits. An amino acid alignment of ASIC3s between 9 relevant rodent species and human identified unique sequence differences that might underlie the proton insensitivity of nmrASIC3. However, introducing nmrASIC3 differences into rat ASIC3 (rASIC3) produced only minor differences in channel function, and replacing nmrASIC3 sequence with that of rASIC3 did not produce a proton-sensitive ion channel. Our observation that nmrASIC3 forms nonfunctional homomers may reflect a further adaptation of the naked mole-rat to living in an environment with high-carbon dioxide levels.

## Introduction

Acid-sensing ion channels (ASICs) are part of the epithelial sodium channel (ENaC)/degenerin (DEG) superfamily of ion channels and are implicated in a diverse range of physiological and pathophysiological processes, ranging from learning and memory to mechanosensation and pain (1). In mammals, there are 4 ASIC encoding genes, which generate 6 distinct ASIC subunits due to splice variants in the *ACCN2* and *ACCN1* genes producing a and b variants of the ASIC1 and ASIC2 subunits respectively: ASIC1a, ASIC1b, ASIC2a, ASIC2b, ASIC3 and ASIC4. The crystal structure of ASIC1 demonstrated that ASICs form trimeric ion channels (2) and although evidence exists for the formation of ASIC/ENaC heteromers (3, 4), it is largely thought that functional ASICs are the result of either homo- or heterotrimeric arrangement of ASIC subunits.

Unlike transient receptor potential vanilloid 1 (TRPV1) that produces a sustained inward current in response to extracellular protons (5), ASICs produce a transient inward current (6). However, being trimeric, the subunit configuration dictates the biophysical characteristics, such as the proton sensitivity for activation, the inactivation time constant and the magnitude of the sustained current in subunit configurations where the current does not completely inactivate in the continued presence of agonist (7). Moreover, the sensitivity to different antagonists is also affected by subunit configuration. For example, the ASIC3 antagonist APETx2 inhibits ASIC3 homomers, as well as heteromers of ASIC3 with ASIC1a, ASIC1b and ASIC2b, but does not inhibit ASIC2a+ASIC3 heteromers (8); APETx2 also relieves hyperalgesia in inflammatory pain models (9, 10). Furthermore, the non-proton ASIC3 agonist 2-Guanidine-4-methylquinazoline (GMQ) causes ASIC3 activation at neutral pH (11), but also modulates the acid sensitivity of other ASIC subunits (12).

Of the 6 ASIC subunits, neither ASIC2b (13) nor ASIC4 (14, 15) form functional homomers, but they are able to form functional heteromers and modulate channel function (7, 13, 16), as can ASIC subunits that have been mutated to make them insensitive to protons as homomers (17, 18). The crystal structure of chicken ASIC1 (cASIC1) identified an acidic pocket region containing three carboxylate pairs (D238–D350, E239–D346 and E220–D408; cASIC1 numbering), which was suggested to be the primary site for proton sensing by ASICs (2). However, ASIC2a lacks D350 and is still functional, whereas ASIC2b also only lacks the D350 carboxylate of the acidic pocket and is not activated by protons (13, 17), results which suggest that regions outside of the acidic pocket must be important for proton activation of ASICs. Indeed, we and others have identified a range of residues on ASIC1a and ASIC2a that when mutated alter proton sensitivity (17–21) and more recently we have shown that the first 87 amino acids of the extracellular domain of rat ASIC2a are required for its proton sensitivity (22).

Understanding the structure-function of ASIC3 is of particular interest because there is substantial evidence supporting involvement of ASIC3 in pain (9, 23–28), as well itch (29), mechanosensation (23, 30) and anxiety (31). Although ASIC3 expression was initially thought to be restricted to the peripheral nervous system (and hence its original name, dorsal root ganglia acid-sensing ion channel, DRASIC (32)), we and others have demonstrated that ASIC3 is also expressed in both the spinal cord and numerous brain regions (33, 34), which makes it even more important to understand how ASIC3 functions because any potential drug that targets ASIC3 for the treatment of pain may also produce side effects with the central nervous system.

Here we investigated the function of ASIC3 cloned from the naked mole-rat, a species that we have previously shown to produce no behavioral response to acid as a noxious stimulus (35). This behavioral indifference to acid is not due to a lack of ASIC-like or TRPV1-like proton-gated currents in sensory neurons, but rather due to an amino acid variation in the voltage-gated Na^+^ channel NaV1.7 that confers enhanced acid block, such that acid acts like an anesthetic, rather than activator of naked mole-rat sensory neurons (36). This is likely a result of adaptation to living in a hypercapnic environment that may induce tissue acidosis (37) and indeed naked mole-rats also show reduced avoidance of ammonia and acid fumes (38, 39), as well as decreased aversion to CO_2_, absence of CO_2_-induced pulmonary edema and enhanced ability to buffer against CO_2_-induced systemic acidosis (40). Although naked mole-rats can more efficiently buffer CO_2_, our previous data regarding the inability of acid to evoke action potentials in an ex vivo preparation (35) that arises from an amino acid variation in NaV1.7 (36) demonstrates that there are likely multiple adaptations to living in a hypercapnic environment. We recently compared ASIC expression between mice and naked mole-rats, finding that whereas ASIC4 is highly abundant in mouse tissues it is the most lowly expressed ASIC transcript in naked mole-rat tissues; the expression pattern of ASIC3 was similar between species (33). At a functional level, we have previously shown that naked mole-rat ASIC1a, ASIC1b and TRPV1 are largely indistinguishable from the mouse orthologs (36, 41) and here we set out to explore the function of naked mole-rat ASIC3 considering its importance to a physiological and pathophysiological processes.

## Results

### Naked mole-rat ASIC3 is insensitive to protons

Primers for cloning mouse ASIC3 (mASIC3) and naked mole-rat ASIC3 (nmrASIC3) were designed based upon the published genome sequences and constructs were made using pIRES2-EGFP or pTarget vectors; rat ASIC3 (rASIC3 in pTracer) was a kind gift from G. Lewin. All constructs were expressed in Chinese hamster ovary (CHO) cells that lack endogenous ASICs (17), and whole-cell patch-clamp electrophysiology was used to measure responses to a pH 4.0 stimulus from a starting pH of pH 7.4. Whereas mASIC3 and rASIC3 robustly responded to protons with a stereotypical transient ASIC-like current (mASIC3: 69 ±13 pA/pF, n = 26, Fig. 1*A*, and rASIC3: 627 ± 92 pA/pF, n = 18, Fig. 1*B*), nmrASIC3 failed to respond with an ASIC-like response to protons, even using a pH 3.0 stimulus, but rather produced a very small, non-inactivating response, similar to that which we have observed previously in non-transfected CHO cells (17) and in CHO cells transfected with the proton-insensitive ASIC2b (22) (pH 4.0, n = 24 and pH 3.0, n = 13, 7 separate transfections, Fig. 1*C*); a summary of all data is given in Table 1. We also investigated whether the non-proton ASIC3 agonist GMQ could activate nmrASIC3, but whereas we observed GMQ-mediated inward currents in cells expressing mASIC3 (Fig. 1*E*, 14.5 ± 3.0 pA/pF, 6 out of 18 pH-sensitive cells responded) and rASIC3 (Fig. 1*F*, 180.6 ± 55.1 pA/pF, n = 8, all pH-sensitive cells responded), nmrASIC3 failed to respond (Fig. 1*G*, n = 11 cells from 4 transfections). One possibility is that nmrASIC3 is retained in the endoplasmic reticulum, as has been proposed for the proton-insensitive mASIC2b, which only gets to the plasma membrane when coexpressed with the proton-sensitive ASIC2a (42). However, using a biotinylation assay to determine plasma membrane expression of mASIC3 and nmrASIC3, transfected either alone, or with rASIC2a, we observed that nmrASIC3 traffics to the plasma membrane, regardless of whether it is transfected alone or with rASIC2a, just like mASIC3 (Fig. 1*H*, a second experiment showed the same plasma membrane trafficking of nmrASIC3). Therefore, the insensitivity of nmrASIC3 to protons and GMQ cannot be explained by a lack of membrane expression.

**Figure 1.**
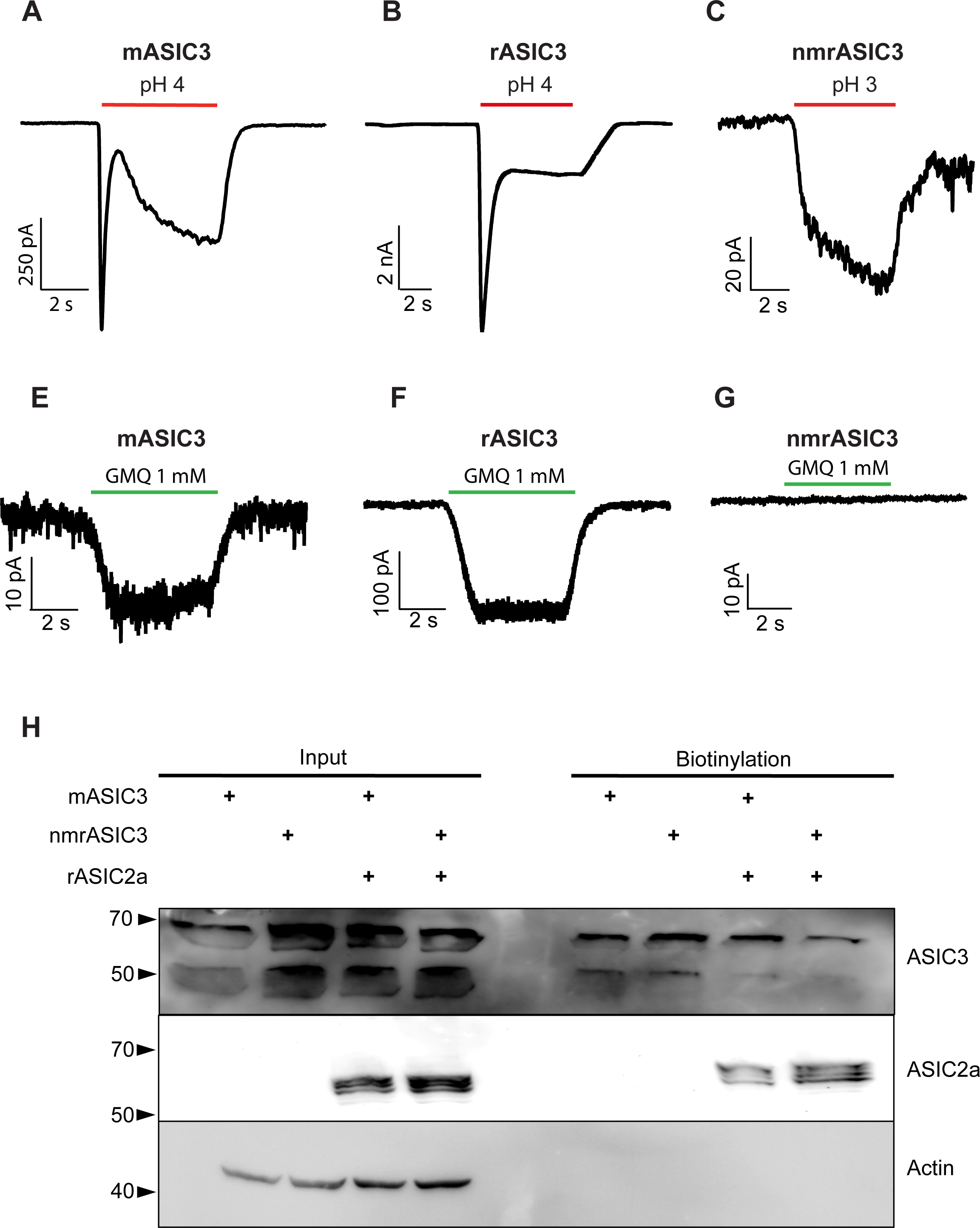
Representative traces of currents recorded during stimulation with low pH and GMQ. Currents recorded from CHO cells expressing mASIC3 (*A*), rASIC3 (*B*) or nmrASIC3 (*C*) stimulated with either pH 4.0 or pH 3.0 solution, and stimulated with 1 mM GMQ (*E-G*). H, Western blot of whole cell lysates (input) and biotinylated surface fraction from cells transfected with mASIC3, nmrASIC3 or cotransfected with m/nmrASIC3 and rASIC2a and stained with anti-ASIC3 antibody, anti-ASIC2 antibody or anti-ß-Actin antibody.

**Table 1.** Summary data of peak current density, I_sus_/I_peak_, EC_50_ and Hill coefficient for all constructs tested. Numbers in parentheses refer to number of cells tested, details of statistical comparisons are not included for clarity, refer to graphs. ND, not determined.

| | Peak current density at pH 4.0 (pA/pF) | $I_{\text{sus}}/I_{\text{peak}}$ at pH 4.0 (%) | Inactivation Time Constant at pH 4.0 (msec) | pH $EC_{50}$ | pH Hill coefficient | Peak current density GMQ (1 mM) (pA/pF) |
| --- | --- | --- | --- | --- | --- | --- |
| nmrASIC1b | 20 ± 3 (21) | 12 ± 2 (17) | 118 ± 7 (21) | 6.02 ± 0.02 (8) | 1.42 ± 0.1 | ND |
| nmrASIC3 + nmrASIC1b | 67 ± 13 (18) | 34 ± 5 (18) | 125 ± 7 (17) | 5.58 ± 0.07 (12) | 0.89 ± 0.09 | ND |
| nmrASIC3 + rASIC2a | 455 ± 217 (10) | 19 ± 4 (6) | 1637 ± 62 | 4.39 ± 0.17 (11) | 1.17 ± 0.16 | ND |
| mASIC3 | 69 ± 13 (26) | 48 ± 4 (23) | 202 ± 12 (23) | 6.01 ± 0.14 (9) | 0.68 ± 0.1 | 14 ± 3 (6) |
| mASIC3-ATG | 109 ± 51 (8) | ND | 352 ± 31 (9) | 6.02 ± 0.14 (10) | 0.65 ± 0.07 | ND |
| mASIC3 + nmrASIC1b | 193 ± 48 (14) | 31 ± 4 (15) | 93 ± 5 (14) | 5.66 ± 0.08 (8) | 0.96 ± 0.09 | ND |
| mASIC3 + rASIC2a | 129 ± 23 (13) | 141 ± 25 (13) | 80 ± 10 (4) | 4.45 ± 0.15 (15) | 0.72 ± 0.06 | ND |
| rASIC3 | 627 ± 92 (18) | 35 ± 5 (18) | 387 ± 52 (12) | 6.39 ± 0.17 (10) | 0.91 ± 0.18 | 181 ± 55 (8) |
| rASIC3A62E | 705 ± 219 (7) | 43 ± 31 (7) | 343 ± 47 (13) | 5.91 ± 0.20 (6) | 0.76 ± 0.07 | ND |
| rASIC3A62E R102H | 673 ± 213 (8) | 26 ± 6 (8) | 191 ± 14 (8) | 6.15 ± 0.13 (7) | 0.72 ± 0.06 | ND |
| rASIC2a | 421 ± 58 (50) | 29 ± 6 (8) | 1220 ± 108 (17) | 4.44 ± 0.08 (14) | 1.62 ± 0.51 | ND |

### nmrASIC3 is functional in dorsal root ganglion neurons

We have previously demonstrated that naked mole-rat dorsal root ganglion (DRG) neurons produce ASIC-like currents in response to acid (36) and that these neurons express nmrASIC3 mRNA (33, 36) and thus we used APETx2, an inhibitor of most ASIC3-containing ASIC trimers, to determine if nmrASIC3 contributes to the acid-sensitivity of these neurons. A pH 5.0 stimulus evoked two types of inward current in naked molerat DRG neurons: rapidly-inactivating, ASIC-like currents and sustained, TRPV1-like currents (Fig. 2*A* and *B*). Exposure to 2 μM APETx2 for 30 seconds caused a significant decrease in the amplitude of the ASIC-like responses evoked by a second pH 5.0 stimulus (61 ± 16 pA/pF vs. 33 ± 10 pA/pF, n = 8, p ≤ 0.05, Fig. 2*A* and *C*), but had not effect upon TRPV1-like responses 5 ± 2 pA/pF vs. 4 ±1 pA/pF, n = 14, Fig. 2*B* and *C*); the inhibition of ASIC-like responses was reversible (47 ±11 pA/pF, p ≤ 0.05). Considering that APETx2 is selective for ASIC3 homomers and most ASIC3 heteromers (8), and that nmrASIC3 does not form proton-sensitive homomers (Fig. 1), these results suggest that nmrASIC3 can form proton-sensitive heteromers with other ASIC subunits, much like the other proton-insensitive ASIC subunits ASIC2b and ASiC4 (13, 16).

**Figure 2.**
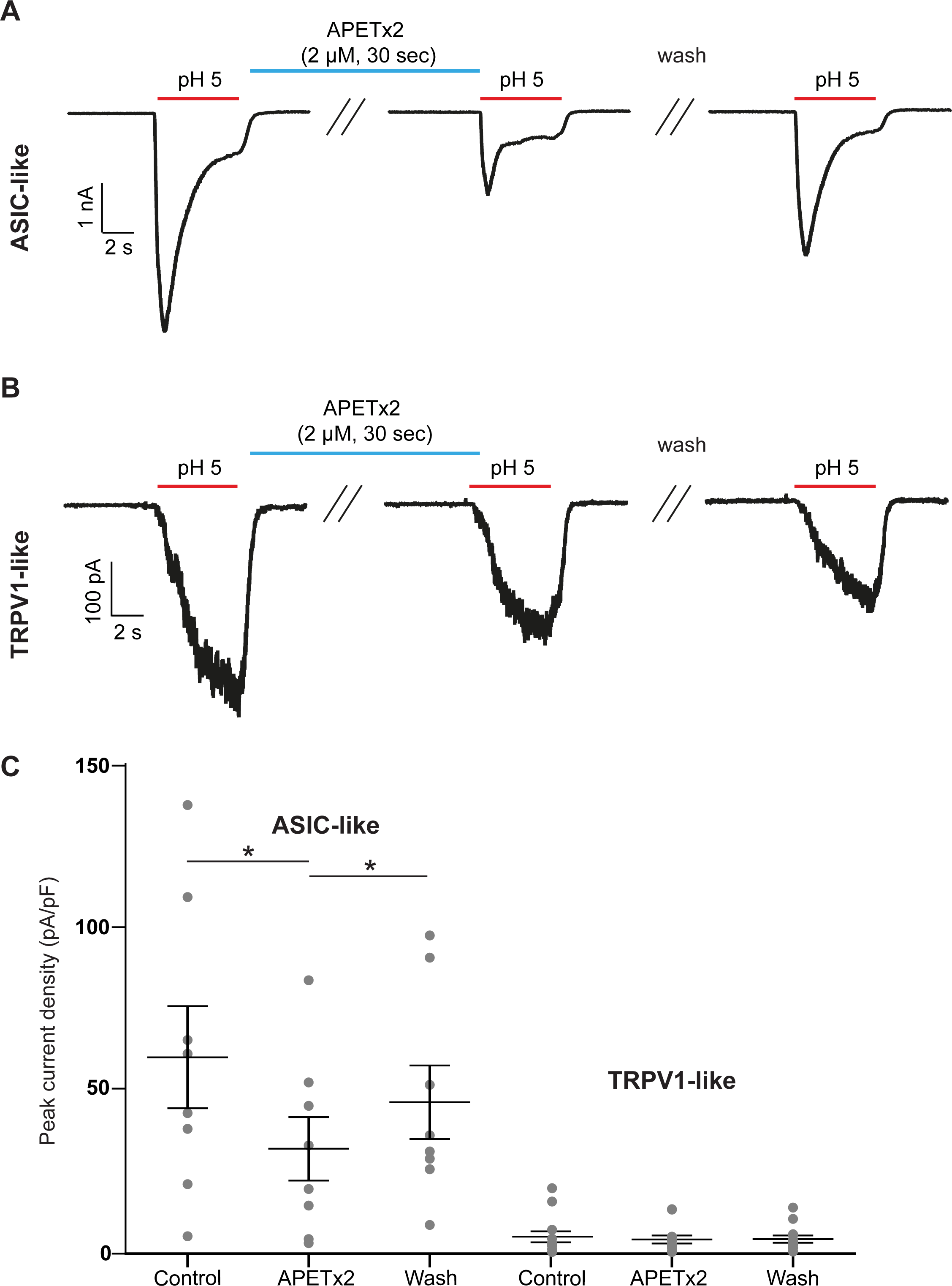
APETx2 blocks transient currents in naked mole-rat DRG neurons. Representative ASIC-like (*A*) or TRPV1-like (*B*) current traces recorded from DRG neurons stimulated with pH 5.0 before APETx2 application, immediately after application of 2 μM APETx2 for 30 s, and after 30 s wash at pH 7.4. *C*, Quantification of results showing that ASIC-like transient currents were significantly and reversibly inhibited by APETx2 while TRPV1-like sustained currents were not affected. Bars represent mean and standard error of the mean. Data were analyzed by paired t-test.**P < 0.01.

### nmrASIC3 forms functional ASIC heteromers with other ASIC subunits

To determine if nmrASIC3 can form proton-sensitive heteromers with other ASIC subunits as DRG neuron data would suggest, we cotransfected either nmrASIC3 or mASIC3 with nmrASIC1b and compared the properties of currents recorded from these cotransfected cells with those only transfected with either mASIC3 or nmrASIC1b. Using a pH 4.0 stimulus, currents recorded from cells transfected with mASIC3 had a peak current density of 69 ± 13 pA/pF (n = 26), an inactivation time constant of 202 ± 12 msec (n = 23) and I_sus_/I_peak_, the sustained current as a percentage of the peak current, was 48 ± 4 % (n = 23, Fig. 3A-D). Currents recorded from cells transfected with nmrASIC1b had a peak current density of 20 ± 3 pA/pF, an inactivation time constant of 118 ± 7 msec (n = 21) and I_sus_/I_peak_ was 12 ± 2 % (n = 17, Fig. 3*A-D*). In cells cotransfected with nmrASIC1b and nmrASIC3, currents were recorded in all instances suggesting that nmrASIC3 does not have a dominant negative effect. Properties of nmrASIC3+nmrASIC1b currents were as follows: peak current density, 67 ± 13 pA/pF (n = 18), inactivation time constant, 125 ± 7 msec (n = 17) and I_sus_/I_peak_ was 34 ± 5 % (n = 18, Fig. 3*A-D*). Similar currents were observed in cells expressing mASIC3+nmrASIC1b: peak current density, 193 ± 48 pA/pF (n =14, p ≤ 0.001 vs. nmrASIC3+nmrASIC1b), inactivation time constant, 93 ± 5 msec (n = 14) and I_sus_/I_peak_ 31 ± 4 % (n = 15, Fig. 3*A-D*). The inactivation time constant of mASIC3 was significantly slower than the inactivation time constants of nmrASIC1b and both mASIC3+nmrASIC1b and nmrASIC3+nmrASIC1b (p ≤ 0.001, Fig. 3*C*) Importantly, for both nmrASIC3+nmrASIC1b and mASIC3+nmrASIC1b mediated currents, the I_sus_/I_peak_ was significantly greater than that of nmrASIC1b homomers (p ≤ 0. 01 and p ≤ 0.05 respectively, Fig. 3*D*; the I_sus_/I_peak_ mASIC3+nmrASIC1b was significantly less than that of mASIC3 homomers, p ≤ 0.05 respectively, but for nmrASIC3+nmrASIC1b there was no significant difference compared to mASIC3 homomers Fig. 3D). These results suggest that both nmrASIC3 and mASIC3 form heteromers with nmrASIC1b to produce currents with a substantial sustained component. Although the large sustained component measured in cells expressing mASIC3 and nmrASIC1b could be the result of measuring a mixture of mASIC3 homomeric currents (large Isus/Ipeak) and nmrASIC1b homomers (small I_sus_/I_peak_) this cannot explain the large I_sus_/I_peak_ measured in cells expressing nmrASIC3 and nmrASIC1b because nmrASIC3 does not form proton-sensitive homomers and thus it is likely that ASIC3 and ASIC1b form heteromers that have a substantial I_sus_/I_peak_ as has been shown by others for rASIC3+rASIC1b (7). A second piece of evidence suggesting that nmrASIC3 can form functional ASIC heteromers is that pH-response curves show that the effective concentration 50 (EC_50_) for mASIC3+nmrASIC1b heteromers was not significantly different from that of nmrASIC3+nmrASIC1b (pH 5.66 ± 0.08, n = 8, vs. pH 5.58 ± 0.07, n = 12, p = 0.91), however, heteromers of nmrASIC3+nmrASIC1b were significantly different from the EC50 of either mASIC3 or nmrASIC1b homomers (mASIC3, pH 6.01 ± 0.14, n = 9 and nmrASIC1b, pH 6.03 ± 0.02, n = 9, p ≤ 0.01 Fig. 3*E*), but not mASIC3+nmrASIC1b (p = 0.053 and 0.05 respectively).

**Figure 3.**
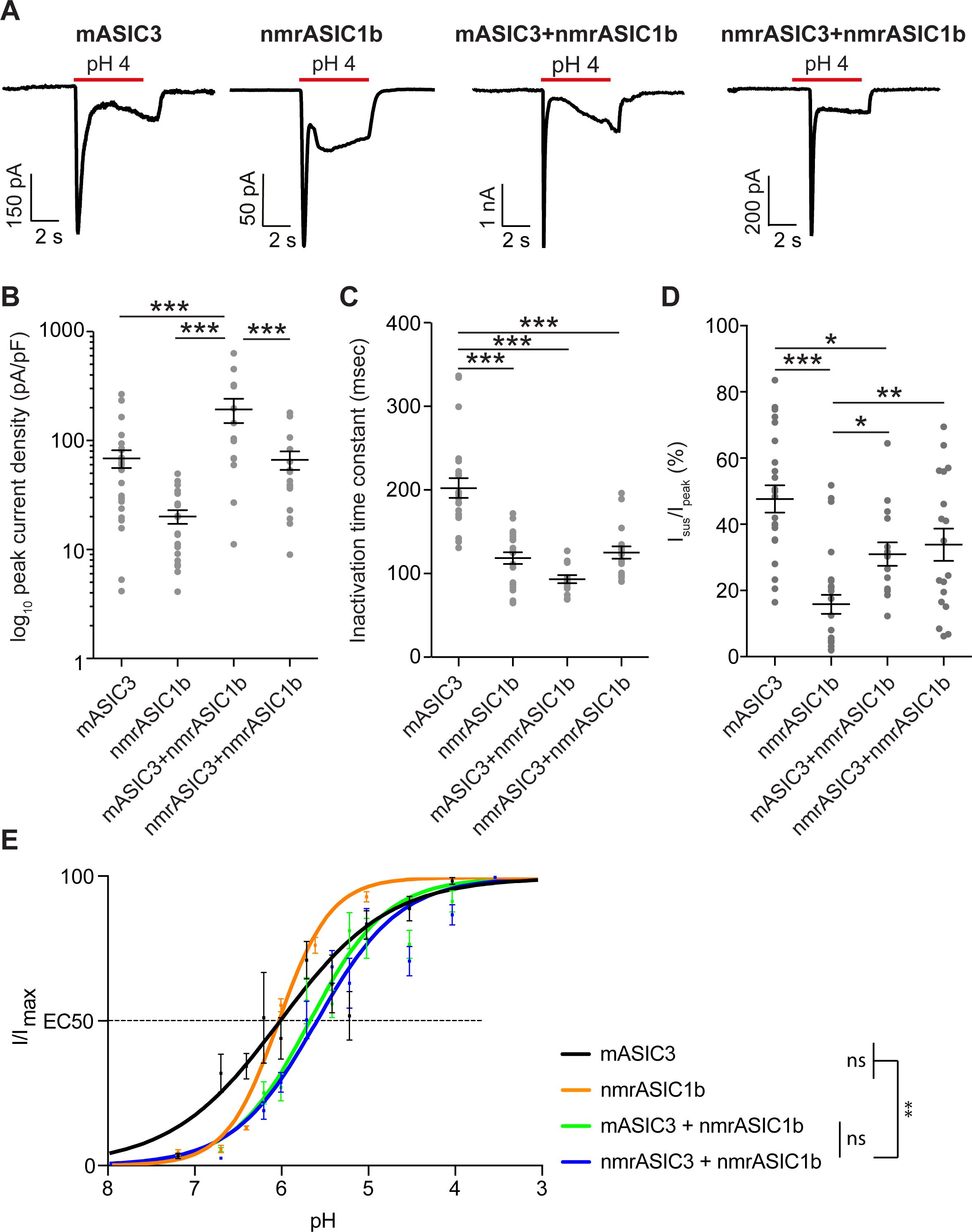
Characterization of CHO cells co-expressing nmrASIClb and mASIC3 or nmrASIC3. *A*, Currents recorded from CHO cells expressing mASIC3, nmrASIClb, mASIC3 + nmrASIClb or nmrASIC3 + nmrASIClb. Quantification of log_l0_ peak current density (*B*), inactivation time constant (*C*) and I_sus_/I_peak_ (*D*). Bars represent mean and standard error of the mean. E, pH-response curves of mASIC3, nmrASIClb, mASIC3 + nmrASIClb and nmrASIC3 + nmrASIClb. Data were analyzed by ANOVA with Tukey’s multiple comparison test.***P < 0.001**P < 0.01 comparing all conditions.

We also investigated the ability of mASIC3 and nmrASIC3 to form heteromers with rASIC2a and found that as shown previously (7,43) coexpression of mASIC3 and rASIC2a resulted in currents with an I_sus_/I_peak_ that was significantly greater than that produced by either mASIC3 or rASIC2a homomers (mASIC3+rASIC2a: 141 ± 25, n = 13 vs. mASIC3: 48 ± 4 %, n = 23, p ≤ 0.001 and rASIC2a 29 ± 6 %, n = 8, p < 0.001 Fig. 4*A* and *D*). By contrast, cells expressing nmrASIC3 and rASIC2a produced currents that did not produce a large I_sus_/I_peak_ (Fig. 4*A* and *D*) and appeared largely indistinguishable from rASIC2a homomers; the lack of a large sustained current in cells expressing nmrASIC3+rASIC2a in response to protons is not necessarily a sign that heteromers are not formed because rASIC2a+rASIC3 heteromers have been reported to have an I_sus_/I_peak_ of ~30 % (28), not to dissimilar to the 19 ± 4 % observed here. Moreover, a small, but significant difference in the inactivation time constant was observed suggesting that heteromers may be formed: nmrASIC3+rASIC2a heteromers inactivated significantly more slowly than mASIC3+rASIC2a heteromers and homomers of either rASIC2a or mASIC3 (nmrASIC3+rASIC2a, 1637 ± 62 msec, n = 9 vs. mASIC3+rASIC2a, 80 ± 10 msec, n = 4, p ≤ 0.001, vs. rASIC2a, 1220 ± 108 msec, n = 17, p < 0.05 and vs. mASIC3, 202 ± 12 msec, n = 23, p ≤ 0.001 Fig. 4*C*). Currents mediated by nmrASIC3+rASIC2a were of significantly greater magnitude than those mediated by mASIC3 (nmrASIC3+rASIC2a 455 ± 217 pA/pF, n = 10, vs. 69 ± 13 pA/pF, n = 26, p ≤ 0.05), whereas those mediated by rASIC2a were significantly greater than those mediated by mASIC3+rASIC2a and mASIC3 (421 ± 58 pA/pF, n = 50, vs. mASIC3+rASIC2a, 129 ± 23 pA/pF, n = 13, p ≤ 0.05, and 69 ± 13 pA/pF, n = 26, p ≤ 0.001 respectively Fig. 4*B*). Examination of pH-response curves showed that both mASIC3+rASIC2a and nmrASIC3+rASIC2a produced currents that were significantly less sensitive to protons than mASIC3, but not significantly different from each other (EC_50_s: mASIC3+rASIC2a, pH 4.45 ± 0.15, n = 15, vs. nmrASIC3+rASIC2a, pH 4.39 ± 0.17, n = 13, p = 0.98, and vs. mASIC3, pH 6.01 ± 0.12, n = 10, p ≤ 0.0001, Fig. 4*E*), and these were also indistinguishable from that of rASIC2a (EC_50_: rASIC2a, pH 4.44 ± 0.05, n = 14, p = 0.99 and p = 0.99 respectively). In summary, based upon biophysical characterization, nmrASIC3 forms functional heteromers with nmrASIC1b (Fig. 3), but the evidence is less clear for heteromeric formation with rASIC2a (Fig. 4) although these two subunits are both present at the plasma membrane when cotransfected (Fig. 1*H*) and the fact that ASIC-like currents are sensitive to inhibition by APETx2 in DRG neurons from naked mole-rats (Fig. 2) strongly supports the premise that although nmrASIC3 produces proton-insensitive homomers it can form functional heteromers *in vivo*.

**Figure 4.**
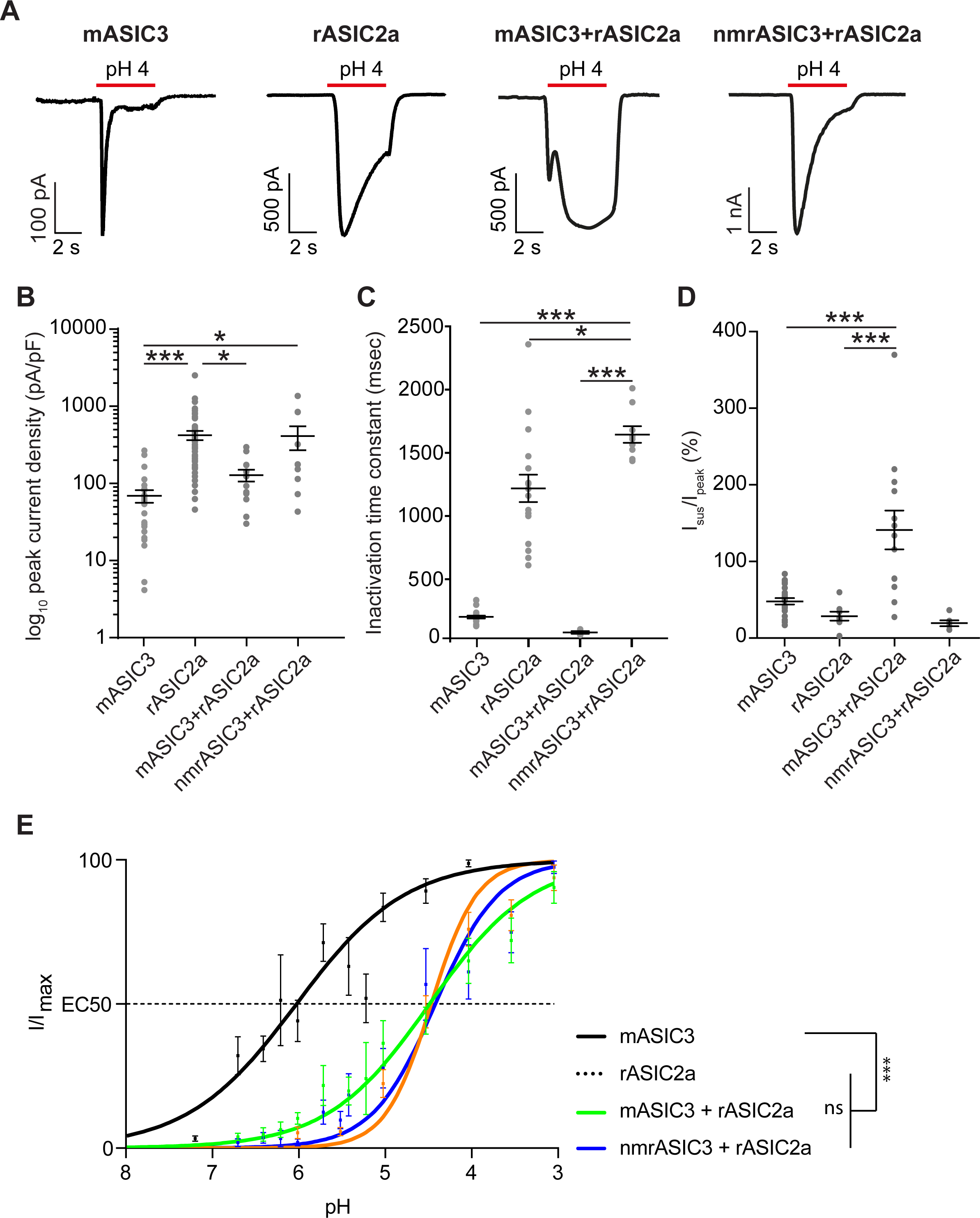
Characterization of CHO cells co-expressing rASIC2a and mASIC3 or nmrASIC3. *A*, Currents recorded from CHO cells expressing mASIC3, rASIC2a, mASIC3 + rASIC2a or nmrASIC3 + rASIC2a. Quantification of logi_0_ peak current density (*B*), inactivation time constant (*C*) and sustained current in proportion to peak current (*D*). Bars represent mean and standard error of the mean. E, pH-response curves of mASIC3, rASIC2a, mASIC3 + rASIC2a and nmrASIC3 + rASIC2a. Data were analyzed by ANOVA with Tukey’s multiple comparison test.***P < 0.001 comparing all conditions.

### Amino acid variations specific to nmrASIC3 do not account for homomeric proton-insensitivity

To determine the molecular basis for proton-insensitivity of nmrASIC3, we aligned the nmrASIC3 amino acid sequence with that of 9 other species (Fig. 5*A-C*). We found only three instances where naked mole-rat residues differed in a region otherwise conserved in all species. Importantly, we included guinea pig, a rodent more closely related to naked mole-rat than mouse or rat, and guinea pig sensory neurones are activated by the non-proton agonist of ASIC3, GMQ demonstrating the functionality of ASIC3 in this species (27). We firstly identified that nmrASIC3 is missing the first methionine of the protein sequence, having instead a methionine at position 7 (Fig. 5*A*). We thus created a version of mASIC3 lacking the initial ATG and further mutated the position 7 leucine for methionine (L7M), termed mASIC3-ATG, and an nmrASIC3 with an added methionine at position 1 and an M7L mutation termed nmrASIC3+ATG. mASIC3-ATG responded to low pH and the EC50 was 6.02 ± 0.14 (n = 10), not significantly different to that of wild-type mASIC3 (p = 0.97, Fig. 5*D*). By contrast, nmrASIC3+ATG, like wildtype nmrASIC3, did not respond to acid. Variation in the initial 7 amino acids of nmrASIC3 cannot therefore explain its proton insensitivity.

**Figure 5.**
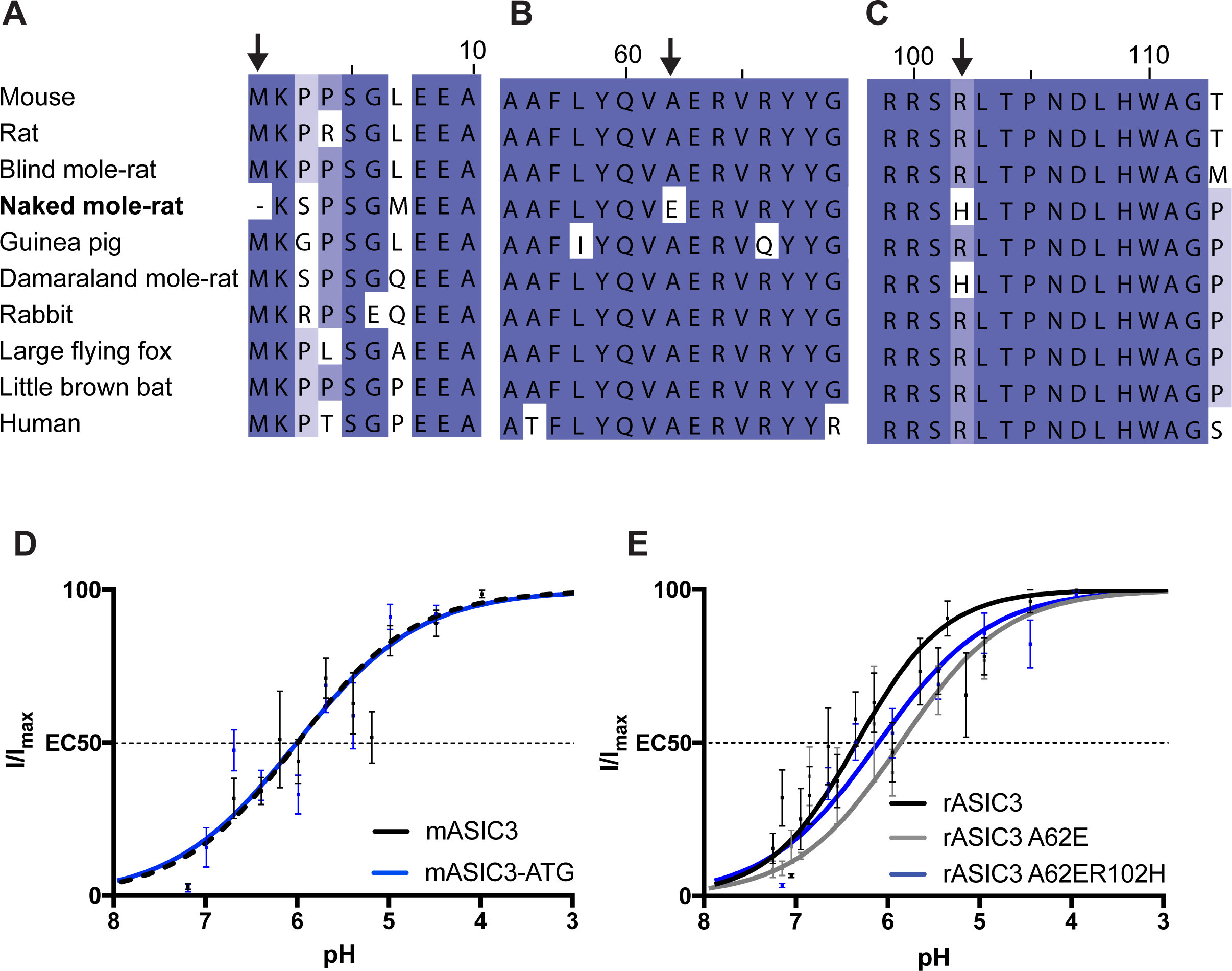
Multiple sequence alignment of rodent and human ASIC3 protein including species closely related to naked mole-rat or living in similarly hypoxic/hypercapnic habitats. *A*, Amino Acids 1 - 10, Methionine is missing in the naked mole-rat (arrow). *B*, Amino acids 55 - 69, change of alanine 62 to glutamate in naked mole-rat (arrow). *C*, Amino Acids 99 - 112, change of arginine to histidine in naked mole-rat and Damaraland mole-rat (arrow). *D*, pH-response curves of mASIC3 and mASIC3-ATG. *E*, pH-response curves of rASIC3, rASIC3A62E, rASIC3A62ER102H. Data were analyzed by T-test.

A second difference exclusive to nmrASIC3 in the comparison made was replacement of alanine at position 62 with glutamate (A62E, Fig. 5*B*), a residue likely within transmembrane domain 1, but close to the start of the extracellular domain (2). Because rASIC3 produced larger currents than mASIC3, we used this construct from this point onwards to determine if mutations altered pH sensitivity. A pH 4.0 stimulus produced inward currents of a similar amplitude in cells expressing either rASIC3A62E or rASIC3 (rASIC3A62E: 705 ± 219 pA/pF, n = 7 and rASIC3: 499 ± 145, n = 10), but currents inactivated significantly more rapidly (rASIC3A62E: 343 ± 47 msec, n = 13 vs. rASIC3: 580 ± 33 msec, n = 5, p ≤ 0.001). The pH-response curve was slightly, but nonsignificantly, shifted to the right (rASIC3A62E: pH 5.91 ± 0.2, n = 6 and rASIC3: 6.39 ± 0.17, n = 10, p = 0.09, Fig. 5*E*).

A third amino acid variation was identified at residue 102, a conserved arginine being replaced in both the naked mole-rat and Damaraland molerat with histidine (Fig. 5*C*) and thus rASIC3A62E was further mutated to produce rASIC3A62ER102H. The pH-response curve for the double mutant rASIC3A62ER102H was not significantly different to that of rASIC3 (pH 6.15 ± 0.13, n = 7, p = 0.33, Fig. 5*E*). As for rASIC3A62E, currents mediated by rASIC3A62ER102H inactivated significantly faster than wild type rASIC3 (rASIC3A62ER102H, 191 ± 14 msec, n = 8, p < 0.001). Thus, it would appear that neither A62 nor R102 are of crucial importance in proton activation of ASIC3 and indeed when expressed in CHO cells neither nmrASIC3E62A nor nmrASIC3H102R resulted in the rescuing of nmrASIC3 proton-sensitivity.

## Discussion

Sensitivity to acid as a noxious stimulus is largely conserved throughout the animal Kingdom (44), but the naked mole-rat is behavioral insensitive to acid (35) due to a variation in NaV1.7, which results in acid anesthetizing, rather than activating, naked mole-rat sensory neurons (36). Moreover, naked mole-rats show behavioral indifference to both ammonia and acid fumes (38, 39), as well as CO_2_ (40) at levels producing avoidance in mice. All of these findings demonstrate likely adaptations to having evolved in a subterranean, hypoxic/hypercapnic environment (45, 46). Here we undertook to investigate the properties of nmrASIC3 because evidence supports a role for ASIC3 in a wide variety of situations, including: pain (9, 23–26), as well itch (29),mechanosensation (23, 30) and anxiety (31). Although nmrASIC3 displays a similar expression profile to mASIC3 (33), we show here that it is insensitive to both protons and the non-proton agonist GMQ when expressed in CHO cells, even though it traffics to the plasma membrane. Considering the 82.7% identity with mASIC3 and 81.6% rASIC3 at the amino acid level, including 95% similarity of the EC domain (EMBOSS matcher algorithm) (47), these findings were unexpected. Much like ASIC2b and ASIC4, which are also insensitive to protons (13, 16), we have produced pharmacological evidence that nmrASIC3 contributes to heteromeric ASIC formation in DRG neurons, such that the ASIC3 subunit containing antagonist APETx2 reversibly inhibits ASIC-like currents, but has no effect on TRPV1-like currents, recorded from naked molerat DRG neurons. It has been well characterized that although homomeric ASIC currents can occur in both peripheral and central neurons (9, 48), it is perhaps more common for ASICs to form heteromers (9, 49) and variation of ASIC subunit configuration has a significant effect upon the sensitivity of channels to protons, their inactivation time constants, and sensitivity to different pharmacological agents (7, 8, 50, 51). As well as the reversible inhibition of proton-gated currents in DRG neurons by APETx2, in a heterologous expression system we observed that nmrASIC3 appears to form heteromers with both ASIC1b and ASIC2a, which would suggest that *in vivo* it acts to modulate ASIC currents, just like ASIC2b and ASIC4.

With regard to understanding the basis of nmrASIC3 proton-insensitivity, manipulation of the few amino acid variations that we identified from a multiple sequence alignment neither abolished rASIC3 proton-sensitivity, nor rescued nmrASIC3 proton-sensitivity and thus it remains unclear why nmrASIC3 fails to respond to protons. Of the 3 carboxylate pairs contained within the acidic pocket identified from the crystal structure of cASIC1 that were proposed to be critical for proton activation of ASICs (D238–D350, E239–D346, and E220–D408; cASIC1a numbering) (2), D238 is replaced by glutamate and D346 is actually replaced by a serine in mASIC3 and rASIC3. Lacking the full set of carboxylates in the proton-sensitive mASIC3 and rASIC3 confirms earlier results from ourselves and others highlighting that although the acidic pocket is involved in proton activation of ASICs, it alone is not responsible for proton activation of ASICs (17–22). In nmrASIC3, E220 becomes D210, E239 becomes D229 and D346 is actually retained D352, thus nmrASIC3 actually has a full set of carboxylates in the acidic pocket, and although two glutamates are replaced by aspartates, which would have a different pKa, they are still protonatable residues. A series of mutations have been made in rASIC3, two of which were similar to those in this study, E63A and R102A, and although EC_50_s for these mutants are not reported, both responded to protons and E63A was observed to slow down the rate of inactivation (52), whereas here we observed that the inactivation time constant was more rapid in rASIC3E62A. However, Cushman *et al.* stimulated using pH 6.0 from a starting point of pH 8.0, whereas here we stimulated using pH 4.0 from a starting point of pH 7.4, which may explain the difference observed. Taken together, from the results presented here it remains unclear why nmrASIC3 is proton-insensitive; there are further nmrASIC3 amino acid variations that remain to be tested, which may yet account for the proton-insensitivity observed, although these are far less species specific than those tested here.

Considering the varied physiology with which ASIC3 is concerned, the fact that nmrASIC3 forms non-functional homomers may be a further adaptation to living in a hypercapnic environment. Future experiments determining the role of ASIC3 in different brain regions of the naked mole-rat will be required to understand just how the proton insensitivity of nmrASIC3 influences brain function.

## Experimental Procedures

### Animals

All experiments were conducted in accordance with the United Kingdom Animal (Scientific Procedures) Act 1986 Amendment Regulations 2012 under a Project License (70/7705) granted to E. St. J. S. by the Home Office; the University of Cambridge Animal Welfare Ethical Review Body also approved procedures. Young adult naked mole-rats were used in this study: 2 males and 1 female aged between 3.5 and 4.5 years. Animals were maintained in a custom-made caging system with conventional mouse/rat cages connected by different lengths of tunnel. Bedding and nesting material were provided along with a running wheel. The room was warmed to 28 °C, with a heat cable to provide extra warmth running under 2-3 cages, and red lighting (08:00 – 16:00) was used.

### Chinese hamster ovary cell culture and transfection

Chinese hamster ovary (CHO) cells (Sigma-Aldrich) were grown using standard procedures in the following medium: Ham’s F-12 Nutrient Mixture (Life Technologies), 10 % fetal bovine serum (Sigma), 1 % Penicillin/Streptomycin 100 U/ml (Life Technologies). 24-hours before transfecting cells, 35 mm dishes (Fisher) were coated with 100 μg/ml poly-L-lysine (Sigma) and cells from a 70-80% confluent flask were trypsinized, resuspended in 5 ml CHO medium and a volume was taken to seed cells at a 1:10 dilution, 2 ml/dish. For transfections, an EGFP expression vector was used to enable identification of transfected cells and DNA was transfected at a ratio of 10:1, ASICx:GFP, or 5:5:1 in c-transfection experiments, using 0.9 μg ASICx DNA and 0.09 μg EGFP DNA (2 μg DNA was used for nmrASIC3); the transfection reagent Lipofectamine LTX (Life Technologies) was used according to the manufacturer’s protocol.

### Cloning and mutagenesis

mASIC3 and nmrASIC3 was amplified from mouse and naked mole-rat whole brain cDNA, respectively, using forward (fw, mASIC3: atgaaacctccctcaggactgga, nmrASIC3: aagagccctcgggatggagga) and reverse primers (rv, mASIC3: ctagagccttgtcacgaggtaaca, nmrASIC3: ctagaatcactagtttgcccgggat), cloned into pIRES or pTarget expression plasmid and confirmed by sequencing. Rat ASIC2a cDNA in a pCI expression plasmid and nmrASIC1b in pEGFP-N3 have been previously described (22, 36). Mutations were inserted with the FastCloning method (53) using primer pairs specific to the construct (mASIC3-ATG fw aaacctccctcaggatggagga,rv cattcctgagggaggttttaccgtcgactgcagaattcga, nmrASIC3+ATG fw, rv aagagcccctcggggctggaggaggctcggagaa, rASIC3A62E fw tctaccaggtggaggagcgggttcg, rv cctggtagaggaaggccgccagcga, rASIC3A62ER102H fw cccactgcgccgctcaca rv gtgaggtgtgagcggcgca, nmrASIC3E62A fw ctaccaggtggctgagcgggtacgcta, rv acctggtagaggaaggctgccagcga, nmrASIC3H102R fw cgctcacgcctcactcccaacga, rv cgtgagcggcgcagcgggttgatgtt).

### Biotinylation

CHO cells were transfected using polyethylenimine (PEI). The cells cultured in 75 cm^2^ flasks were approximately 75% confluent. For transfection, to 1 mL of serum-free DMEM media, 25 μg of total plasmid DNA encoding either mASIC3, nmrASIC3 or rASIC2a was mixed with 15 μL of 7.5 mM PEI. When a combination of plasmids was to be transfected, the concentration of plasmids was split equally. This transfection mixture was incubated for 10 minutes at room temperature and added drop-by-drop to the flask which was replaced with fresh growth media prior to addition of the transfection mixture. For isolation of cell-surface biotinylated proteins, 48-hours post transfection, the growth medium was removed from cells. Ice-cold HEPES buffer saline (HBS) (140 mM NaCl, 1.5 mM Na_2_HPO4.2H_2_O, 50 mM HEPES, pH 7.05) containing 0.2 mg/mL biotin-sulfo-NHS (Thermo Fisher Scientific, Cat. # 21331) was added to cells and incubation was carried out for 60 minutes on ice. Subsequently, biotin containing HBS was removed and cells were washed at least 3 times with 15 mL of tris-buffered-saline (25 mM Tris-HCl, 150 mM NaCl & 10 mM EDTA, pH 7.4). Cells were collected in the same buffer, pelleted at 1000 g for 5 minutes at 4C. The pellet was solubilized in solubilization buffer (25 mM Tris-HCl, 150 mM NaCl, 10 mM EDTA, 1% Triton X-100 and 1 mg/mL protease inhibitor (Roche)) for 60 minutes at 4 °C with continuous mixing. This lysis mixture was centrifuged at 50,000 g for 60 minutes at 4 °C and the supernatant was incubated with 50 μ L of monomeric avidin-coated agarose beads (Thermo Fisher Scientific) for 2-hours at 4 °C with continuous mixing. The protein-bead complexes were collected by centrifugation at 20,800xg for 10 minutes, washed with the solubilization buffer at least 3 times with a mixing time of 5 minutes between washes. The protein was eluted from the beads using 50 μ L Laemmli buffer for immunoblotting. For electrophoresis, 20 μ L of the protein-laemmli buffer mix was loaded in the lanes of a 10% acrylamide and SDS-PAGE was carried out. Proteins were then transferred onto a polyvinyl difluoride (PVDF) membrane, blocked with 5% milk-TBS tween20 solution for 60 minutes at room temperature, probed with primary antibody at 4 °C overnight (anti-ASIC2, Abcam, ab77384 (1:250), and anti-ASIC3, Boster, PA1938 (1:250)), washed with 5% milk-TBS tween20 solution and incubated with secondary antibody (anti-mouse HRP, used at 1:1000, Thermo Fisher Scientific, Cat. # 31430; anti-rabbit HRP, used at 1:1000, Bio-Rad, Cat. # 1706515) for 2-hours at room temperature. Blots were washed in distilled water and then developed with West Pico Chemiluminescent Substrate (Thermo Fisher Scientific). Protein samples from beads were checked for any ‘biotin-permeabilization’ by probing for actin (negative control, A2228, 1:500, Sigma Aldrich).

### Electrophysiology

Dorsal root ganglion (DRG) neurons were cultured as described previously (36, 41). Whole-cell patch clamp recordings from CHO cells were performed at room temperature 24-hours after transfection and recordings from DRG neurons 24-hours after culturing. For all ASIC experiments, the intracellular solution contained 110 mM KCl, 10 mM NaCl, 1 mM MgCl_2_, 1 mM EGTA, 10 mM HEPES, 2 mM Na_2_ATP, 0.5 mM Na_2_GTP in MilliQ water; pH was set to pH 7.3 by adding KOH and the osmolality was adjusted to 310-315 mOsm with sucrose. The extracellular solution contained 140 mM NaCl, 4 mM KCl, 2 mM CaCl_2_, 1 mM MgCl_2_, 10 mM HEPES (solutions > pH 6) or MES (solutions < pH 6), 4 mM Glucose in MilliQ water; osmolality was adjusted to 300-310 mOsm with sucrose and pH was adjusted with NaOH and HCl; unless stated otherwise, e.g. for test pH solutions, the extracellular solution was pH 7.4. Patch pipettes were pulled from glass capillaries (Hilgenberg) using a Model P-97, Flaming/Brown puller (Sutter Instruments) and had a resistance of 4-10 MΩ. Data were acquired using an EPC10 amplifier (HEKA) and Patchmaster software (HEKA). 2 μM APETx2 were added to the pH 7.4 solution for DRG neuron experiments and solutions containing the synthetic ASIC3 agonist GMQ (Sigma) in pH 7.4 were diluted from a stock solution of 50 mM dissolved in DMSO. For measurement of current amplitude and inactivation time constant, a protocol of 5 s of pH 7.4 followed by a 5 s stimulus with pH 3 or pH 4 then a return to pH 7.4 solution for 5 s was used; the holding potential was -60 mV for both DRG neurons and CHO cells. For pH-response curves, 2.5 s stimuli between pH 3 and pH 6 (ASIC2) or 1s stimuli between pH 4 and pH 7.2 (ASIC1, ASIC3) were applied in random order with 30 s of bath solution in between stimuli to minimize desensitization, although this is not a prominent feature of ASIC1b, ASIC2a or ASIC3 (22, 54).

### Data analysis

Statistical analysis was performed in Prism (Graphpad), which was also used to plot data. Peak current density was analyzed by measuring the size of the peak current compared to the baseline current (average current measured over 4 seconds prior to stimulation). The absolute current size was then divided by capacitance of the cell to result in normalized peak current density (pA/pF). Peak current density data was transformed using yi = log_10_(xi). The inactivation time constant τ was measured using a built-in function of Fitmaster. Statistical analysis was performed in GraphPad Prism using repeated measures analysis of variance (ANOVA) with Tukey’s multiple comparisons for DRG neuron data or ordinary one-way analysis of variance (ANOVA) and Tukey’s multiple comparisons, comparing data from each construct with every other construct of the same experiment for transformed peak current density data and inactivation data of CHO cell experiments. Results are expressed as mean and standard error of the mean (SEM), unless otherwise stated; this might, however, not necessarily represent the statistical differences for peak current density data, which were transformed as above. Sustained current to transient current ratio was calculated by measuring the size of the sustained current at the end of the stimulus compared to the baseline current and dividing his value by the peak current (I_sus_/I_peak_ x 100). For pH-response curves, all measurements were transformed to percent of the maximum peak current (I/I_max_ x 100) for each cell. The EC_50_ and Hill coefficient were determined by plotting individual pH-response curves for every cell using Graphpad Prism. Outliers were identified using Prism’s ROUT method with Q = 1% and those cells were eliminated from the dataset. The mean EC_50_ and Hill coefficient for each condition were then used to calculate an average curve using the Hill equation, which was plotted along with the mean and SEM of measured values. Figures were made using Adobe Illustrator.

### Multiple sequence alignment

ASIC3 protein sequences were obtained from the NCBI genome database or ENSEMBL (mouse NM_183000,naked mole-rat PREHGLG00000022115, rat NM_173135.1, guinea pig ENSCPOT00000020999, blind molerat XM_008847215.1, Damaraland mole-rat XM_010637570.1, rabbit ENSRNOG00000058308,human ENSG00000213199, large flying fox ENSPVAG00000007091, little brown bat ENSMLUG00000028175). Naked mole-rat sequences that were not previously annotated were identified using NCBI’s basic local alignment search tool (BLAST) online. Sequences were aligned using MAFFT version 7 using default settings with an un-alignment factor of 0.8. Sequences were visualized and manipulated in Jalview. Alignments are shaded by BLOSUM62 score which indicates the likelihood that two amino acids are aligned because they are homologous (55). Dark blue indicates that the residues agree with the consensus sequence (>80% homology), medium blue that they have a positive BLOSUM62 score (corresponding to >62% likelihood of homology), light blue represents >40% homology and white residues are not homologous (<40% homology).

## Acknowledgments

L.-N. S. was a student in the Biotechnology and Biological Sciences Research Council Doctoral Training Programme [Grant BB/J014540/1] and a recipient of the David James Pharmacology Award. S. S. was supported by the Cambridge International and European Trust. E. St. J. S. acknowledges a Royal Society Research Grant (RG130110). G.C. and E. St. J. S. acknowledge funding from Arthritis Research UK (Grant Reference 20930). All authors thank members of the Smith lab for assistance in the lab, especially J. Bartlett, J. Raby and L. Rockall, and are grateful to the support of animal technicians in particular H. R. Forest and A. J. Robinson. The authors are also grateful for discussion of the project with Prof. G Lewin and Dr D. Omerbašić.

## Author Contributions

LNS and EStJS conceived and designed the study. LNS created constructs, conducted electrophysiology experiments and analyzed the data. SS conducted biotinylation experiments. GC conducted some CHO cell electrophysiology experiments and analyzed the data. EStJS conducted DRG neuron electrophysiology experiments and analyzed the data. All authors contributed to writing the manuscript.

